# The structural basis of drugs targeting protein-protein interactions uncovered with the protein-ligand interaction profiler PLIP

**DOI:** 10.1101/2025.03.04.641378

**Authors:** Sarah Naomi Bolz, Philipp Schake, Celina Stitz, Michael Schroeder

**Affiliations:** Biotechnology Center (BIOTEC), CMCB, Technische Universität Dresden, Dresden, Germany; Center for Scalable Data Analytics and Artificial Intelligence (ScaDS.AI), TU Dresden, Dresden, Germany

## Abstract

Promiscuity of drugs and targets plays an important role in drug-target prediction, ranging from the explanation of side effects to their exploitation in drug repositioning. A specific form of promiscuity concerns drugs, which interfere with protein-protein interactions. With the rising importance of such drugs in drug discovery and with the large-scale availability of structural data, the question arises on the structural basis of this form of promiscuity and the commonalities of the underlying protein-ligand (PLI) and protein-protein interactions (PPI).

To this end, we employ the protein-ligand interaction profiler, PLIP, to characterize the PLI and PPI of MDM2/p53, Bcl-2/BAX, XIAP/Casp9, CCR5/gp120, and BRD/H4. We show that the wealth of existing complexes in PDB, of predicted protein structures, and of molecular docking results gives deep insights into the design principles for drugs targeting protein interactions.

Drugs targeting protein interaction interfaces are a promising avenue in drug discovery. Understanding drug and target promiscuity at a structural level will pave the way to deliver on this promise.

## 1. Introduction

### Promiscuity of drugs and targets in drug discovery

Drugs that are able to interact with multiple different targets or targets that can accommodate multiple different drugs are known as promiscuous. In drug discovery, promiscuity is an essential concept because it can influence both the therapeutic potential and side effects of a compound (37489516, 23524195). While the interaction with multiple targets may cause unwanted side effects (22722194), promiscuity can also lead to polypharmacology, where the interaction with more than one target enhances therapeutic effects (24946140). Moreover, promiscuity provides a molecular basis for drug repositioning approaches, which aim to use already established drugs for new indications (30310233, 32419905). A drug capable of binding to two targets associated with different conditions may be effective in treating both (32419905). Drug repositioning has the potential to reduce drug development time and cost since approved drugs have known safety profiles and are already in production (30310233). During the COVID-19 pandemic, drug repositioning proved particularly valuable as it accelerated the availability of treatments (32889701, 32870481).

### Drugs targeting protein-protein interactions

The inhibition of protein-protein interactions (PPIs) represents a specific form of promiscuity. It is based on the premise that the target interacts not only with its natural protein binding partner but also with a drug. PPIs play essential roles in numerous biological processes, making them attractive targets for therapeutic intervention. In many cases of PPI inhibition, the drug binds directly to the protein-protein interface, with the protein-ligand interactions (PLIs) mimicking key aspects of the PPIs (26119925). Understanding and characterization of therapeutically important PPIs can serve as a rationale for the discovery of such drugs (26119925).

Protein structure data are of high value in this context because they offer insights into the interactions in atomic detail (37489516). The protein data bank (PDB) (36420884) currently stores experimentally determined structures of about 500k (partially redundant) protein-protein interfaces and about 2200 drug-like molecules bound to these interfaces (38907989).

### Characteristics of drug-target and domain-domain interfaces

In general, protein-protein interfaces are characterized by large, flat surfaces that lack well-defined pockets, making them challenging drug targets (32968059). Despite these common features, protein-protein interfaces largely vary in their binding stabilities, involving both stable and transient interactions that enable a wide range of protein functions (27050677, 18355092). Moreover, interfaces can be categorized into distinct structural classes that require different drug discovery approaches (27050677). Critical for the understanding of protein-protein interfaces is also the finding that interface residues do not equally contribute to binding, but a specific set of residues, known as hot spots, accounts for most of the binding energy (17546660). Identifying these hot spots can facilitate the detection of suitable binding sites and guide drug discovery (24997383).

### Analysing interfaces and binding sites

While experimentally determined structures are of high quality, the availability of these data is limited. Protein structure prediction tools, such as AlphaFold, can now generate protein monomer structures with high accuracy (34265844). The AlphaFold Database currently contains over 200 million protein structure predictions, massively expanding the amount of structural data compared to the PDB (34791371). Moreover, recent progress has resulted in precise models of PLI and PPI (38718835), which could facilitate large-scale screenings for PPI inhibitors.

Tools that analyse the interactions in PLI and PPI structures in detail are essential to developing virtual screening approaches for PPI inhibitors. One such tool is the Protein–Ligand Interaction Profiler (PLIP) (33950214). PLIP automatically detects and visualizes non-covalent interactions in protein-ligand complexes, providing insights into the binding modes and key residues involved. The latest update of PLIP introduced protein-protein interactions. PLIP identifies eight different interaction types: hydrophobic interactions, hydrogen bonds, water bridges, π-stacking, π-cation interactions, halogen bonds, salt bridges, and metal complexation. In virtual drug screenings, PLIP has been successfully used to identify drug repositioning candidates based on similar PLI patterns (26873186,37280244,32459810,34285770,27626687). Identifying commonalities between PPI and PLI patterns could, therefore, be a promising approach to screen for drugs targeting PPI.

### Clinical success of drugs targeting protein-protein interactions

Although PPIs are challenging targets, they hold great promise for drug discovery because they are fundamental to many biological processes. Several drugs modulating PPIs have recently entered clinical trials. Lu et al. summarized the successes of these modulators in a 2020 review article (32968059). They identified 16 small molecules targeting PPIs that were being studied in clinical trials at that time. Given the importance and promise of drugs targeting PPI and the availability of structural models of proteins, we set out to shed light on the structural basis of these drugs. We will illustrate five of the examples mentioned above on a structural basis using PLIP to compare similarities and differences between drug-target and protein-protein interaction interfaces.

## 2. Results

### PPI interfaces are large and flat

Protein-protein interfaces are considered more challenging to target than traditional ligand binding sites (29908451). In contrast to ligand binding sites, which are often deep pockets in enzymes or G-protein coupled receptors, protein-protein interfaces are typically large and flat (15060526, 32968059). Structural analyses of protein-protein complexes in the 1990s found that a standard-sized interface buries a surface area of 1,200 to 2,000 Å² (2204619, 8552589, 9925793). Large interfaces can reach sizes of 4500 Å² (9925793) and often consist of multiple recognition patches (11948787).

### PPI interfaces are diverse

Furthermore, rather than being a uniform group of drug targets, protein-protein interfaces have diverse properties. The character of a protein-protein interaction can be obligate (permanent) or non-obligate (transient), which translates into a wide range of observed binding affinities, from picomolar to high micromolar (27050677). Obligate protein-protein interfaces are larger and more hydrophobic than non-obligate interfaces, possibly reflecting the influence of the hydrophobic effect on binding stability (27050677, 18355092). Furthermore, interfaces can be categorized into structural classes based on whether they involve interactions between globular domains or peptide segments and on whether these elements undergo conformational changes upon binding (27050677, 16524830). Since the interacting structural elements define the shape of the binding site, the choice of an appropriate drug discovery approach depends on the structural interface class (27050677). Different interface types vary in their druggability and present distinct challenges in drug discovery (27050677, 35562538).

### PPI interface hot spots

A key concept in the understanding of protein-protein interactions and the identification of druggable surface regions at interfaces are hot spots (17546660, 24997383). It originates from the finding that critical residues of the interface contribute disproportionately to the binding energy (7529940, 9653027). Notably, hot spots do not spread over the whole interface but are arranged in clusters, so-called hot regions (15644221). Hot spots can be experimentally identified by alanine scanning mutagenesis (24997383). This strategy is complemented by multiple computational hot spot prediction methods (24997383). For instance, early prediction methods used simple energy models to predict hot spots based on shape complementarity and non-covalent interactions (12381794, 14634848). The identification of hot spots can guide drug discovery approaches for protein-protein interfaces by highlighting sites where small molecules may bind with high affinity (24997383). Interface hot spots are enriched in Trp, Tyr, and Arg residues. These residues can contribute to multiple types of favorable non-covalent interactions (17546660). Both Trp and Tyr can participate in π-interactions, hydrophobic contacts, and hydrogen bonds. Arg may be involved in a salt bridge and hydrogen bonds (17546660). Hot spot residues and the interactions they engage in can be analysed with PLIP.

### PPI and PLI interfaces in PLIP

We wanted to quantify whether the non-covalent PLI and PPI differ in terms of the quantity and quality of these interactions. Thus, we selected 134,743 protein structures with a small molecule bound from the PDB and ran a PLIP analysis. We found that hydrogen bonds, hydrophobic contacts, water bridges, and salt bridges are the most abundant interactions with 37, 28, 11, 10%, followed by metal complexation, π-stacking, π-cation interactions, and halogen bonds at 9, 3, 1, 0.2%. For the 948,941 hydrogen bonds in PDB, distances average to 2.51 Å and angles to 141 degrees, which is in line with literature (14696187). For a detailed break-down, see Table 1. Similarly, we composed a set of interacting proteins by selecting from the SCOP2 database (31724711) 453 PDB entries with exactly two chains, A and B. We analysed the interacting chains with PLIP and found - consistent with PLI - that in PPI hydrogen bonds, hydrophobic contacts, water bridges, and salt bridges are the most abundant interactions with 33, 40, 18, 8 %, followed by π-stacking, π-cation interactions at 0.6% each. However, in contrast to PLI, there were no metal complexations and halogen bonds. The distances and angles of the 6,087 hydrogen bonds in PPI average to 2.42 Å and 144 degrees. As mentioned earlier, PPIs are larger than PLIs, and this is also reflected in the number of interactions that PLIP detects. The average number of PLIP interactions for PLIs is 12, whereas it is 48 for PPIs. With these high-level statistics on molecular interactions between PPI and PLI in hand, let us now turn to concrete examples of drugs that target protein-protein interaction interfaces.

**Table 1:** Statistics of non-covalent interactions detected by PLIP in PDB structures of protein-ligand and protein-protein interfaces.

|  |  | hydrogen bond | hydrophobic | water bridge | salt bridge | metal complexation | pi-stacking | pi-cation | halogen bond |
| --- | --- | --- | --- | --- | --- | --- | --- | --- | --- |
| protein-ligand | % of interactions | 37 | 28 | 11 | 10 | 8.9 | 3.0 | 0.97 | 0.23 |
|  | ∅ length | 2.51 | 3.71 | 6.66 | 4.48 | 2.20 | 4.29 | 4.24 | 3.37 |
|  | ∅ angle | 141 | - | 135 | - | - | 39.6 | - | 115 |
| protein-protein | % of interactions | 33 | 40 | 18 | 7.8 | 0 | 0.63 | 0.56 | 0 |
|  | ∅ length | 2.42 | 3.73 | 6.46 | 4.20 | - | 4.68 | 4.22 | - |
|  | ∅ angle | 144 | - | 139 | - | - | 56.3 | - | - |

### Drugs targeting protein-protein interactions

Insights into the design principles of protein-protein interactions are vital to discovering and developing small molecules targeting these interactions. Analysis of the non-covalent contacts, as detected by PLIP, provides molecular details of these principles. From success stories of protein-protein interaction inhibition, we can learn how molecular insight from interaction analyses can facilitate drug discovery and how to face inherent challenges.

Lu et al. have listed 16 successful small-molecule inhibitors of protein-protein interaction that have entered clinical trials (32968059). These drugs target eight different protein-protein interfaces. We screened the PDB to identify the cases for which structural data of the protein-protein inhibition are available. Remarkably, for all but one protein-protein interface, there is at least one experimentally determined 3D structure of the complex in the PDB. Eight drugs targeting five different interfaces have both protein-protein and protein-ligand complex structures available in the PDB (Table 2). They comprise the interactions of cancer target MDM2 with p53, cancer target Bcl-2 with BAX, inhibitor of apoptosis XIAP with apoptotic enzyme caspase-9, chemokine receptor CCR5 with the HIV spike protein gp120, and bromodomain-containing proteins with acetylated histones.

**Table 2:**
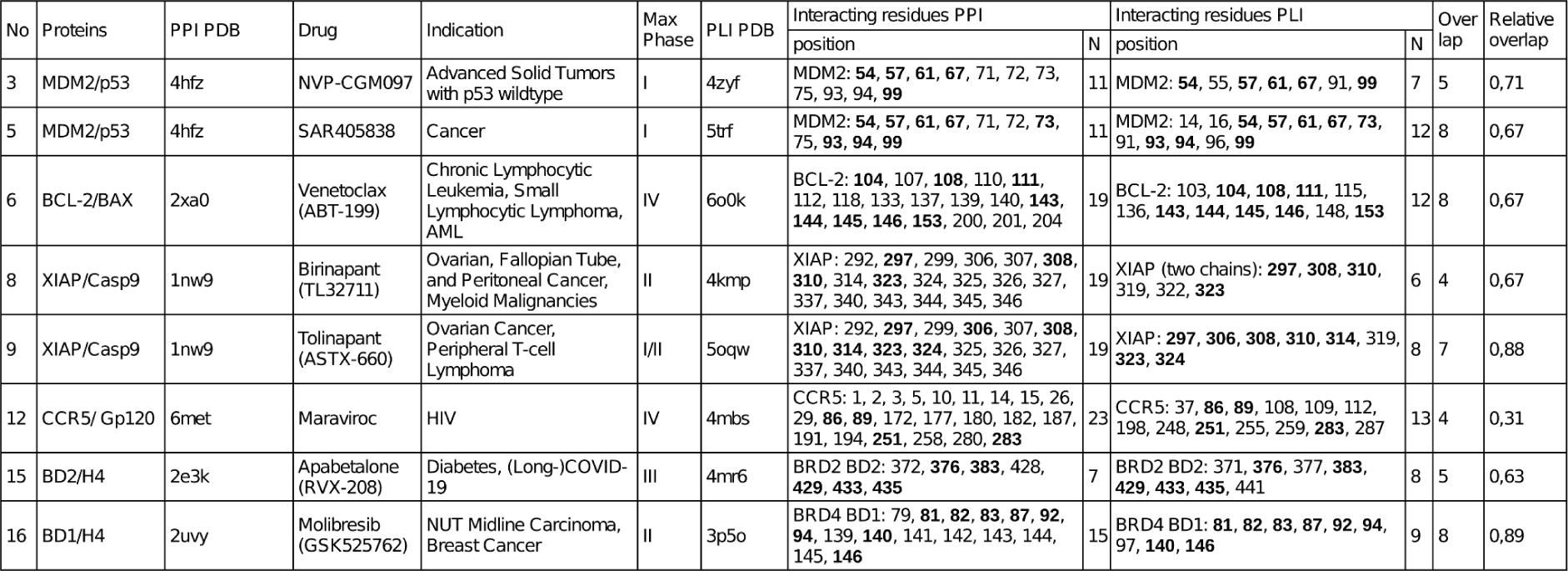
Examples of drugs in clinical trials targeting protein-protein interactions taken from Lu et al. (32968059). No indicates the number in Lu et al. Contacted residues detected by PLIP are given, with overlaps between PPI and PLI in bold.

### PLIP interactions for drugs targeting PPI

For these cases, PLIP detected between 7 and 23 contacting residues at protein-protein interfaces and between 7 and 13 at protein-ligand interfaces, which is consistent with PPI being larger than PLI. While six protein-protein interfaces have more contact residues than their corresponding protein-ligand interfaces, two drugs interact with one more residue than in the respective protein-protein interfaces. For instance, the HIV spike protein gp120 contacts 23 and the HIV drug maraviroc 13 residues of CCR5, respectively. On the other hand, an acetylated histone peptide forms non-covalent interactions with 7 residues of the second bromodomain of bromodomain-containing protein (BRD)2, whereas apabetalone establishes interactions with 8 residues of this domain.

Notably, the contacted residues show considerable overlap between the protein-ligand and the protein-protein interfaces in all cases. The proportion of the residues that are contacted in both interfaces relative to the residues contacted by the drug ranges from 30% to 89%. For example, of 12 Bcl-2 residues contacted by venetoclax, 8 are also contacted by the BH3 domain of BAX. The smallest overlap shows maraviroc, with 4 contacted residues also being involved in the protein-protein interaction.

These observations confirm the finding that protein-protein interfaces are usually larger than protein-ligand interfaces (27050677, 29908451). However, they also point out that in successful protein interaction inhibition by small molecules, the protein-protein interfaces can be as small as protein-ligand interfaces, potentially making them more easily to target than very large interfaces. Protein-protein interfaces between a globular domain and a peptide segment, like MDM2/p53 or Bcl-2/BAX, are considered more druggable than other types of interfaces (27050677). Remarkably, all inhibitors bind their targets in a promiscuous manner, as the protein-protein and protein-ligand interfaces show overlaps in the binding site residues.

To gain a clearer understanding of the binding patterns underlying protein interaction inhibition, we examined the non-covalent PLIP interactions involved in inhibiting the protein-protein interfaces with available structural data. For a summary of the following examples, see Table 3.

**Table 3:**
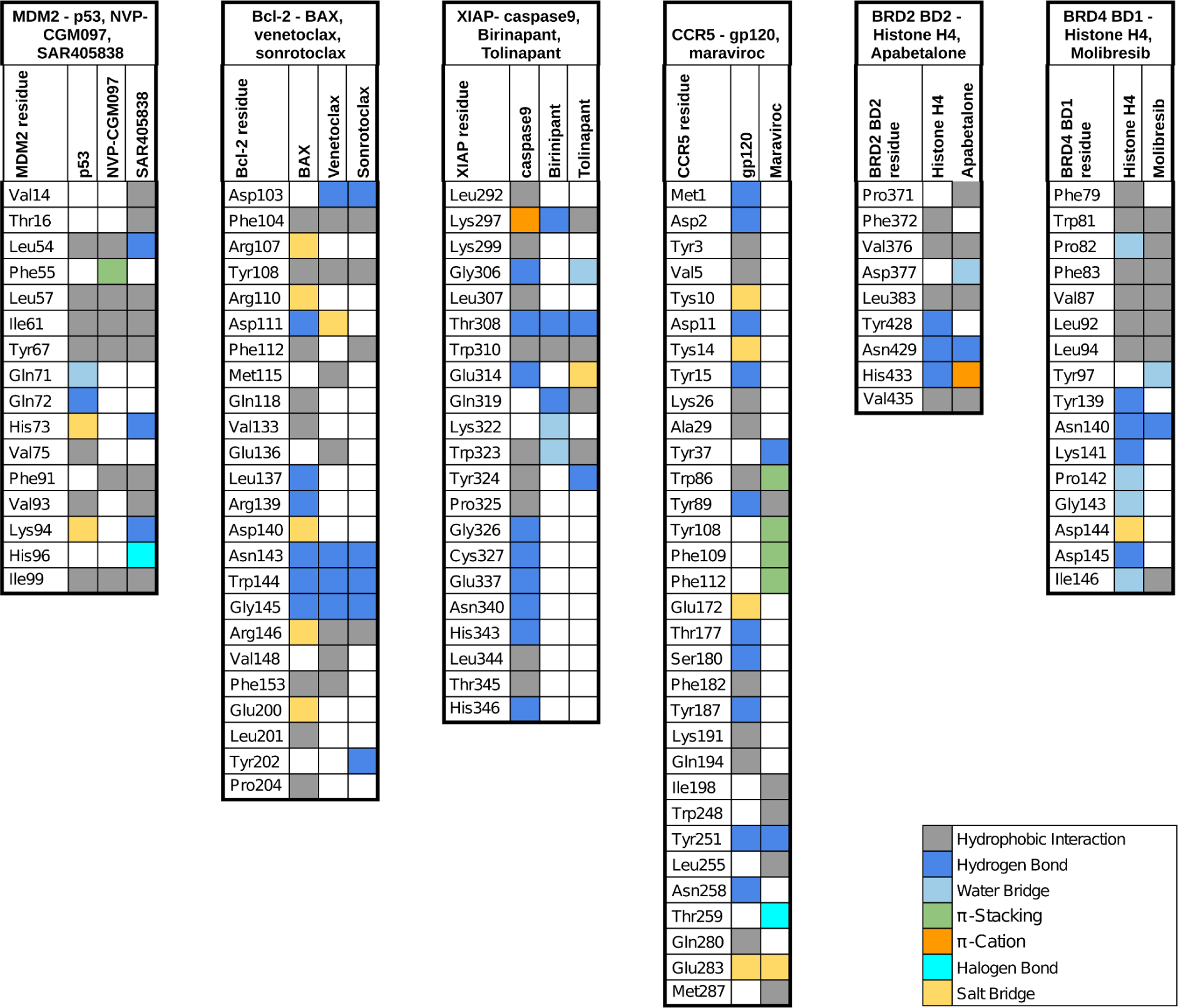
Non-covalent PLIP interaction types (grey: hydrophobic interaction, blue: hydrogen bond, light blue: water bridge, yellow: salt bridge, green: π-stacking, orange: π-cation, cyan: halogen bond). If a residue participated in multiple interaction types, one of them was selected.

### MDM2/p53: biological background

The E3 ubiquitin ligase mouse double minute 2 (MDM2) is a key regulator of the tumor suppressor p53 (28886379). Upon activation by stress or DNA damage, p53 induces cell cycle arrest, DNA repair, or apoptosis if the damage is severe (28886379). In unstressed cells, however, p53 levels are kept low to enable the normal growth and development of cells (28886379). MDM2 forms a negative feedback loop with p53 to maintain homeostasis (19776744). While p53 enhances MDM2 expression, MDM2 binds to the transactivation domain of p53, inhibiting its ability to transactivate MDM2 and other downstream genes (19776744). MDM2 also promotes p53 degradation via ubiquitination and facilitates its translocation to the cytoplasm (28886379, 11713287). Mutations in p53 frequently occur in cancer, almost always leading to a loss of function (35361963). In some human cancers with wild-type p53, MDM2 overexpression presents an alternative mechanism of p53 inactivation (27194168). In these cases, restoring p53 tumor suppressor activity by inhibiting its interaction with MDM2 is a promising therapeutic strategy (27194168). Several compounds targeting the MDM2/p53 interaction have already entered clinical trials (32651541, 35831864). Nutlins, the first potent and selective MDM2 inhibitors, entered clinical trials in the 2000s. Since then, alternative scaffolds have emerged to target MDM2. For instance, SAR405838 features a spirooxindole scaffold designed to mimic the interaction of the p53 residue Trp23 with MDM2, while NVP-CGM097 possesses a dihydroisoquinolinone scaffold identified through virtual screening (35831864).

### MDM2/p53: interactions

The MDM2 inhibitors block the MDM2/p53 interaction by targeting the binding site of the p53 transactivation domain. The co-crystal structure of MDM2 and a fragment of the p53 transactivation domain has revealed a deep hydrophobic cleft in MDM2 that engages with the p53 residues Phe19, Trp23, and Leu26 (8875929, 23897459). PLIP detects extensive hydrophobic interactions between this triad and the MDM2 residues Leu54, Leu57, Ile61, Tyr67, Val75, Val93, and Ile99 (see Figure 1). In total, PLIP finds 17 interactions at the MDM2/p53 interface.

**Figure 1:**
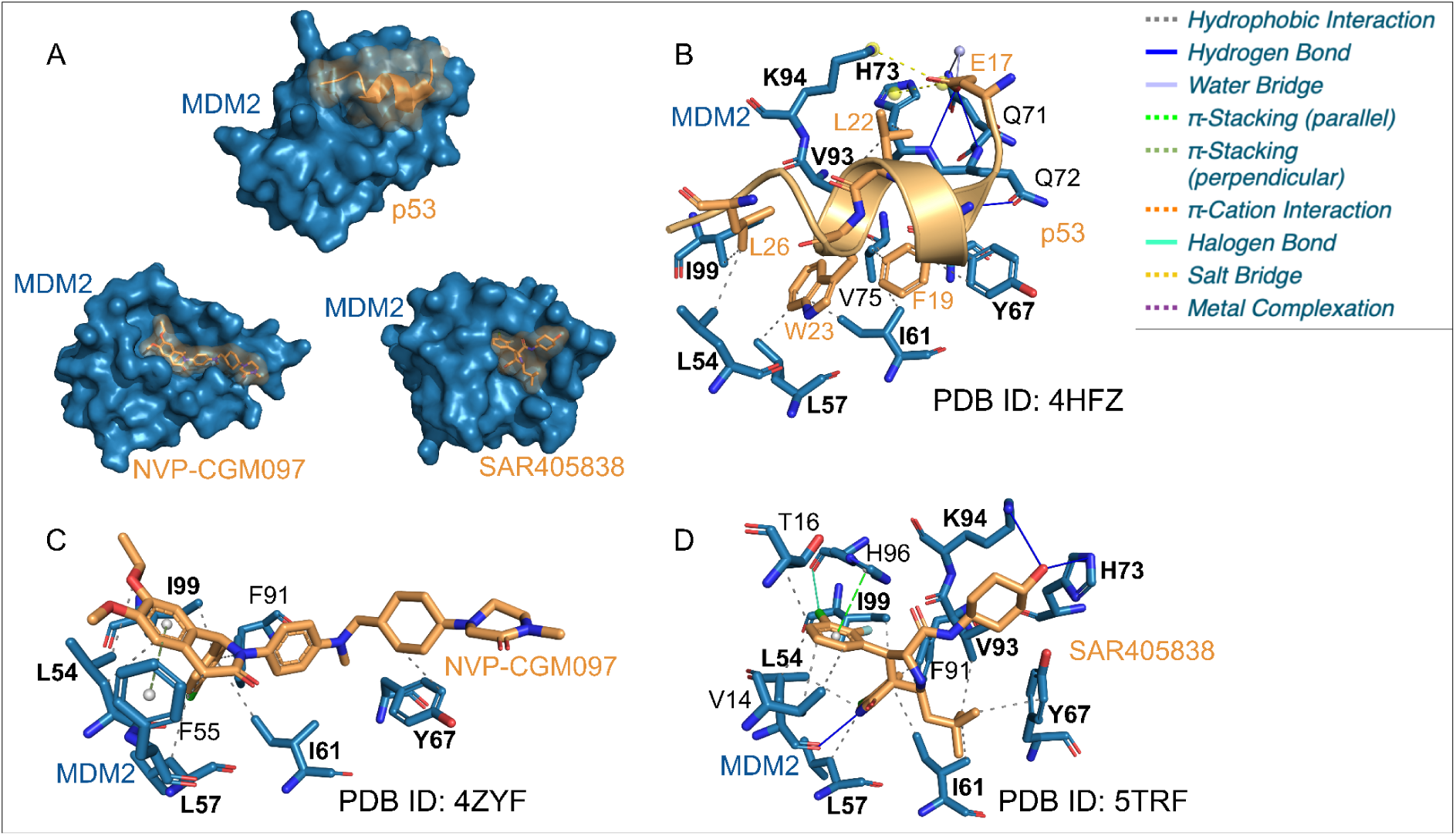
Non-covalent interactions at the interface of MDM2 and p53 or the two inhibitors NVP-CGM097 and SAR405838. The interaction patterns of the inhibitors overlap with the pattern of p53. **A** Surface representation of the complex structures of the interactions of MDM2 (blue) with p53, NVP-CGM097, and SAR405838 (orange). **B** Non-covalent interactions detected by PLIP between MDM2 (blue) and the p53 transactivation domain (orange). p53 residues F19, W23, and L26 are critical for binding and PLIP finds extensive hydrophobic interactions with these residues. **C, D** PLIP interactions of the inhibitors NVP-CGM097 (C, orange) and SAR405838 (D, orange) with MDM2 (blue) agree well with the pattern between MDM2 and p53 (overlapping residues in bold), but involve the additional interaction types π-stacking and halogen bonds.

The MDM2 inhibitors NVP-CGM097 and SAR405838 bind the cleft in a very similar way. Overall, they form 9 and 18 interactions with MDM2, respectively. Both compounds contact the binding site residues Leu54, Leu57, Ile61, Tyr67, and Ile99 via hydrophobic interactions. In particular, the observed interactions with SAR405838 confirm that the spirooxindole scaffold mimics the hydrophobic contacts of the p53 residue Trp23 with the MDM2 residues Leu57 and Ile61. SAR405838 also binds to Val93.

Along with the hydrophobic interactions of the triad, the p53 residue Glu17 forms multiple polar interactions with MDM2, involving hydrogen bonds, salt bridges, and a water bridge with the binding site residues Gln71, Gln72, His73, and Lys94. While NVP-CGM097 does not contact these residues, SAR405838 establishes hydrogen bonds with His73 and Lys94. The interaction of NVP-CGM097 and SAR405838 is further supported by π-stacking interactions with Phe55 and His96, respectively.

Interestingly, SAR405838 contributes a halogen bond to the interaction. Halogen bonds are rarely observed in natural protein-protein interactions, as the 20 standard amino acids lack halogen atoms. The binding mode of SAR405838 indicates that small molecules can use a broader set of interaction types for binding.

### Bcl-2/BAX: biological background

Bcl-2 is a pro-survival protein that sequesters BH3 domain-containing pro-apoptotic proteins of the Bcl-2 family like BAX and BIM (9704409, 21060336). These proteins are responsible for mitochondrial outer membrane permeabilization, a critical process for apoptosis in vertebrates (18314333). Suppression of apoptosis by Bcl-2 contributes to cancer cell survival, and Bcl-2 overexpression is frequently observed in various cancers (21415859). Therefore, small-molecule inhibitors that mimic pro-apoptotic BH3 domains hold great promise as therapeutic agents (25952548, 23291630, 38211332). Venetoclax is a potent and selective Bcl-2 inhibitor that was approved for Chronic Lymphocytic Leukemia (CLL) in April 2016 (23291630). Yet, patients frequently develop resistance against the drug during therapy through a mutation (G101V) at the venetoclax binding site (31160589).

### Bcl-2/BAX: interactions

Bcl-2 sequesters pro-apoptotic Bcl-2 family proteins by directly binding their alpha-helical BH3 domain. A co-crystal structure with the BAX BH3 domain (21060336) shows BH3 binding in a cavity formed by four α-helices of Bcl-2. PLIP analysis identifies 29 non-covalent interactions at the interface, involving several hydrophobic interactions between the BH3 domain and Bcl-2 residues Phe104, Arg107, Tyr108, Phe112, Val133, Leu137, Phe153, Leu201, and Pro204 (see Figure 2). However, the interaction also has a strong hydrophilic profile, including salt bridges with Bcl-2 residues Arg107, Arg110, Asp140, Arg146, and Glu200, as well as hydrogen bonds with Arg107, Asp111, Leu137, Arg139, Asn143, Trp144, Gly145, and Arg146.

**Figure 2:**
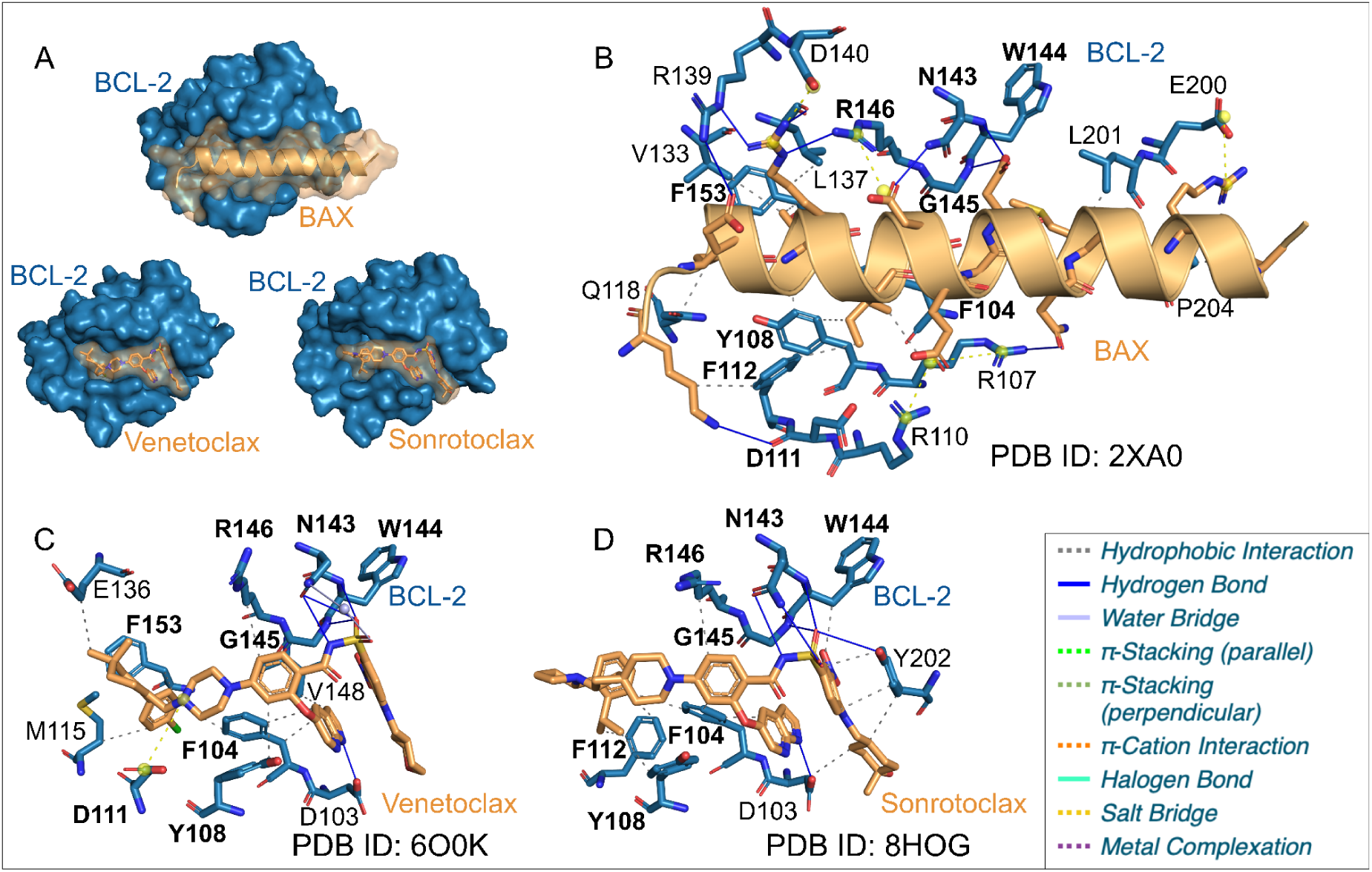
Non-covalent interaction patterns detected by PLIP of Bcl-2 with the BH3 domain of BAX vs the approved drug venetoclax and the more recent inhibitor sonrotoclax. Both inhibitors show significant overlaps (residues in bold) with the Bcl-2/BAX interface. **A** Surface representation of the complex structures of Bcl-2 (blue) with the BH3 domain of BAX, venetoclax, and sonrotoclax (orange). **B** Non-covalent interactions between Bcl-2 (blue) and the BAX BH3 domain (orange). **C, D** Interaction patterns of Bcl-2 (blue) with venetoclax (C, orange) and sonrotoclax (D, orange).

The two Bcl-2 inhibitors venetoclax and sonrotoclax (see Figure 2) bind to the same site as the BH3 domain and, in total, engage in 16 and 18 non-covalent interactions with Bcl-2, respectively. Both compounds contact the BH3 binding site residues Phe104, Tyr108, Asn143, Trp144, Gly145, and Arg146. However, their binding modes differ slightly in the interactions of venetoclax’s chlorophenyl group and sonrotoclax’s isopropylphenyl group with residues of the BH3 binding site. Venetoclax forms a salt bridge with Asp111 and a hydrophobic interaction with Phe153, while sonrotoclax establishes a hydrophobic interaction with Phe112. Interestingly, venetoclax induces a conformational shift of Phe112, but sonrotoclax keeps the residue in its BH3-bound state. It has been shown that sonrotoclax overcomes venetoclax resistance (38211332). The observed differences in the binding mode of sonrotoclax explain its ability to escape resistance mutations (38211332).

Despite the differences, all complexes show hydrophilic interactions with a tetrad of Bcl-2 residues - Asn143, Trp144, Gly145, and Arg146 - along with hydrophobic contacts with Phe104 and Tyr108, suggesting that these interactions are crucial for binding. Such comparisons of interaction patterns could help identify key interactions in protein-protein interfaces, providing valuable insights for the discovery of new drugs.

### XIAP/Casp9: biological background

Caspase-9 functions as an initiator enzyme within the intrinsic apoptotic pathway, which cleaves downstream effector caspases and leads to the apoptosis of the cell (12620238). The enzyme in its monomeric form is inactive and requires homodimerization for activation (11734640, 12620238). X-linked inhibitor of apoptosis protein (XIAP), a key member of the inhibitor of apoptosis protein (IAP) family, can inhibit the dimerization process of caspase-9 by binding to caspase-9 with its third baculoviral IAP repeat (BIR3) domain and therefore traps caspase-9 in an inactive monomeric state.

Since apoptosis is essential for maintaining cellular homeostasis and its dysregulation is a key characteristic of cancer, XIAP overexpression has been identified in various types of cancer (22293567, 24092992). Therefore, research on Smacs (Second

Mitochondria-Derived Activator of Caspases), which are endogenous XIAP antagonists, and Smac mimetics has opened new therapeutic avenues for targeting XIAP in cancer therapy (22293567).

### XIAP/Casp9: interactions

The BIR3 domain of XIAP serves as a heterodimerization interface for caspase-9 while also acting as a binding site for Smac and Smac mimetics such as tolinapant and birinapant. The co-crystallized structure of XIAP-BIR3 with the catalytic domain of caspase-9 reveals an extensive hydrophobic protein–protein interface. The interface is stabilized by a total of 26 interactions, as detected by PLIP, comprising a combination of hydrophobic interactions and hydrogen bonds. Three major interaction motifs contribute to this interface: a helix of the XIAP residues 337–346, which engages in pronounced hydrophobic interactions and hydrogen bonding, a flexible loop of XIAP consisting of Trp223, TYR224, Pro225, Gly226, and Cys227, which also participates in hydrophobic interactions and hydrogen bonding, and a shallow groove of XIAP formed by residues 292–314 that accommodates the N-terminal region of caspase-9 (see Figure 3). This groove is primarily stabilized by hydrophobic interactions and hydrogen bonds but also includes a distinct π-cation interaction at Lys297 (12620238).

**Figure 3:**
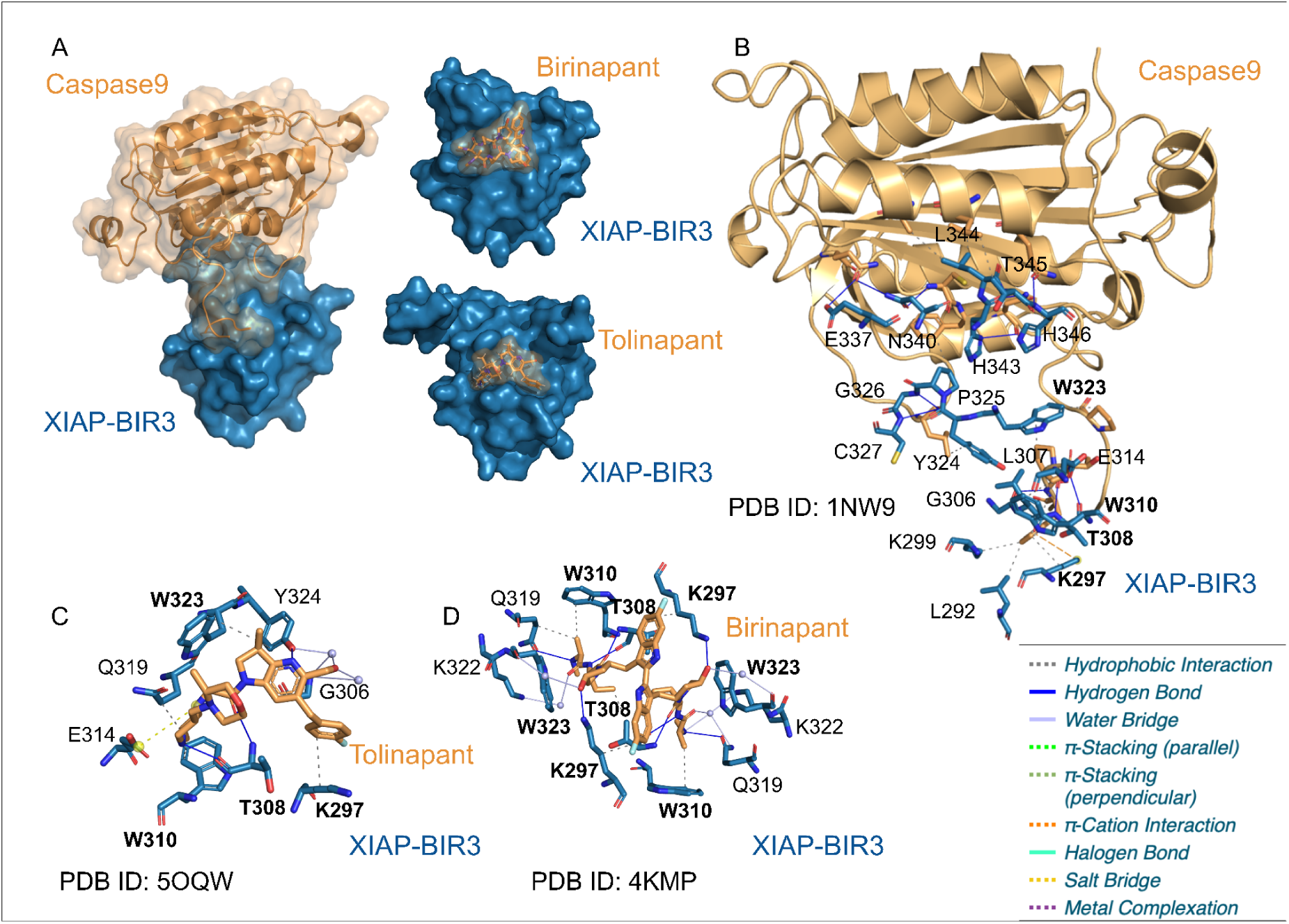
Non-covalent interaction patterns detected by PLIP of XIAP with caspase-9 or the inhibitor birinapant and tolinapant. The interaction between caspase-9 and XIAP forms a large interface, whereas the interaction between XIAP and the inhibitors involves a smaller interface, with a significant overlap observed between the interaction of both inhibitors, which is also recognizable in the XIAP/caspase-9 interaction. **A** surface representation of the complexe structures of XIAP (blue) with caspase-9, birinapant, tolinapant (orange). **B** non-covalent interactions detected by PLIP between XIAP (blue) and caspase-9 (orange). **C,D** PLIP interaction patterns between XIAP (blue) and the inhibitors tolinapant (C, orange) and birinapant (D, orange).

Compared to the XIAP/caspase-9 interaction, the binding of Smac mimetics tolinapant and birinapant to XIAP forms smaller interfaces, with 12 and 19 (involving two chains of XIAP) interactions, respectively. Neither of the two inhibitors binds to the helix comprising residues 337–346. Instead, they interact exclusively with the loop and the shallow groove, as well as additional residues that were not detected in the caspase-9/XIAP interaction by PLIP. However, both inhibitors similarly interact with XIAP and exhibit substantial overlap in the residues involved in binding.

A common feature across all three complexes is the presence of hydrophobic interactions and two hydrogen bonds at Thr308, as well as at least one hydrophobic interaction with Trp310. Differences in the interaction types can be seen at residue Lys297, where the XIAP/caspase-9 complex exhibits a hydrophobic interaction as well as a π-cation interaction. In the XIAP/tolinapant complex and the XIAP/birinapant complex, there are no π-cation interactions, but rather a hydrophobic interaction and, in the case of birnipant, also a hydrogen bond.

Additionally, interactions with Gln319 are exclusive to Smac mimetic complexes, with tolinapant forming a hydrophobic interaction, whereas birinapant engages in both a hydrogen bond and a water bridge.

These results highlight that the interaction of caspase-9 and XIAP forms a larger interface than the interaction between XIAP and the inhibitors tolinapant and birinapant.

### CCR5/gp120: biological background

The human immunodeficiency virus (HIV) infects human cells of the immune system, leading to progressive immune deficiency and, if untreated, acquired immunodeficiency syndrome (AIDS) (8093551, 30629090). The viral spike, which is composed of three gp120-gp41 heterodimers, initiates human cell entry by engaging with host receptors (31262533). Infection first requires gp120 to bind the primary CD4 receptor, followed by interaction with a co-receptor, e.g., chemokine receptors CCR5 or CXCR4 (31262533). Subsequent dissociation of the gp120-gp41 complex and refolding events in gp41 allow the virus to fuse viral and human cell membranes (31262533). Blocking viral entry is one of the strategies used in HIV combination therapy (17617275, 18691983). Maraviroc, which was approved in 2007, directly binds to CCR5, thereby inhibiting the interaction between gp120 and the co-receptor (16251317, 18691983, 24030490).

### CCR5/gp120: interactions

The interface between gp120 and CCR5 consists of two major binding sites (30542158). The first site is formed by the extended N-terminal segment of CCR5 and the bridging sheet of gp120, which becomes accessible only after CD4 binding (30542158). The second binding site involves a loop of gp120 that inserts deeply into a pocket formed by the seven transmembrane helices of CCR5 (30542158). In total, the two binding sites participate in 45 non-covalent interactions. The interactions in the first binding site are governed by several hydrogen bonds and salt bridges with the sulfated tyrosine residues Tyr10 and Tyr14 of CCR5 (see Figure 4). This aligns with the observation that tyrosine sulfation contributes to HIV entry (10089882). In the second binding site, PLIP detects a network of hydrogen bonds between the gp120 loop and CCR5 residues Tyr89, Glu172, Thr177, Ser180, Tyr187, Tyr251, and Asn258. The interaction also involves two salt bridges, where Arg304 and Arg313 in the gp120 loop engage with Glu172 and Glu283 in CCR5, respectively. Hydrophobic contacts with CCR5 residues Leu26, Ala29, Trp86, Glu172, Thr177, Phe182, Lys191, Gln194, and Gln280 further contribute to binding.

**Figure 4:**
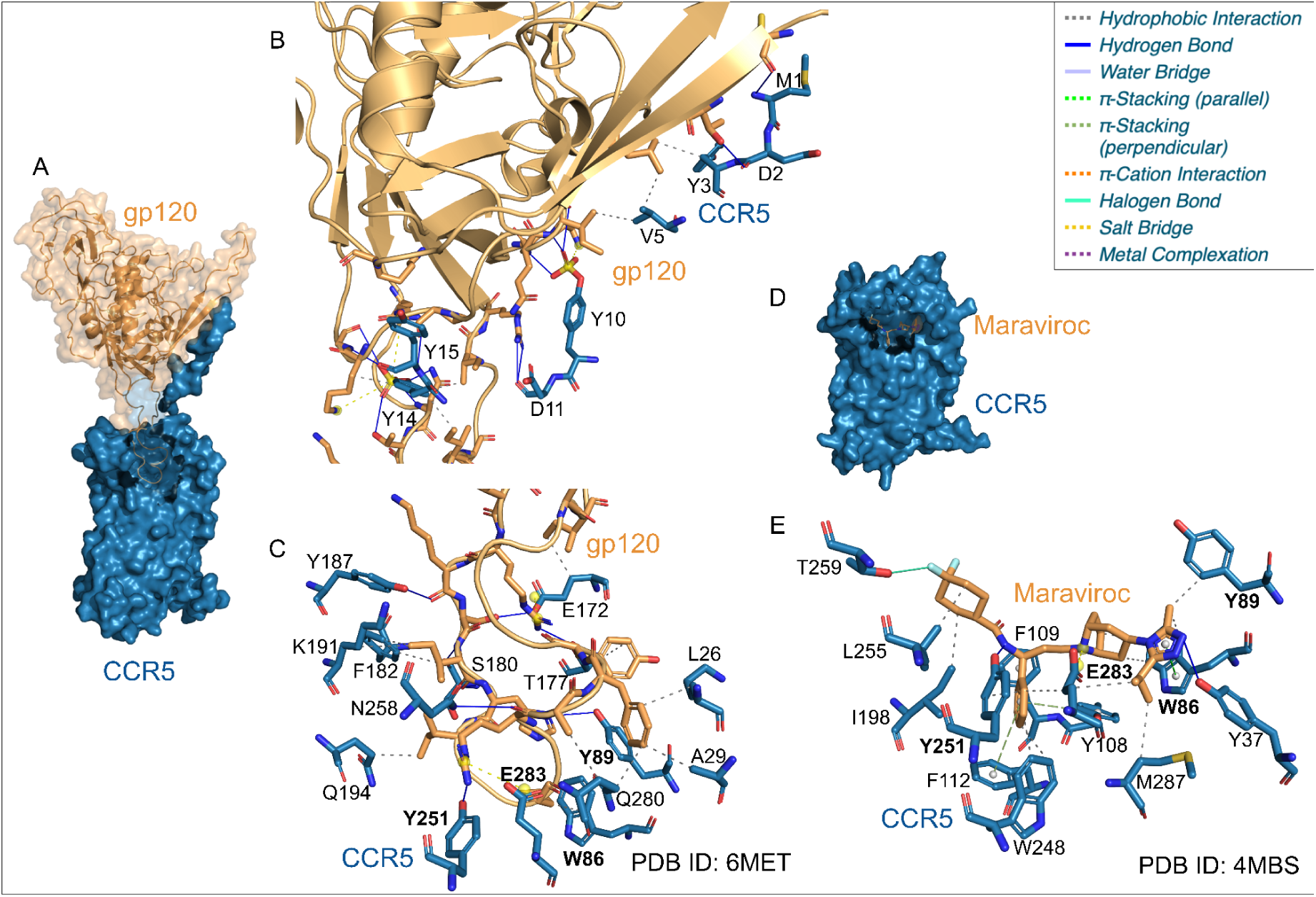
PLIP interaction patterns of CCR5 with the HIV spike protein gp120 vs the approved drug maraviroc. The large CCR5/gp120 interface is divided into two parts, of which Maraviroc only binds one. Overlaps in the contacted residues are bold. **A** Surface representation of the CCR5/gp120 interface. Part of CCR5 (blue) are transparent to make the gp120 (orange) loop visible. **B** Non-covalent interactions in the first binding site of the CCR5/gp120 interface involving the extended CCR5 N-terminal (blue) and the gp120 bridging sheet (orange). **C** Second binding site of the CCR5/gp120 interface. A gp120 loop (orange) interacts with the transmembrane helices of CCR5 (blue). **D** Surface representation of a complex structure of CCR5 (blue) with the approved inhibitor maraviroc (orange), partially transparent to show the position of the drug. **E** Interaction pattern of CCR5 (blue) with maraviroc (orange).

Maraviroc binds to the second binding site of CCR5 via 20 non-covalent interactions, directly competing with the gp120 loop for interaction (24030490, 30542158). Like the gp120 loop, the drug engages with residues Trp86, Tyr89, Tyr251, and Glu283. However, it also contacts residues of CCR5 located deeper within the cavity, specifically Tyr108, Phe112, Trp248, and Met287. As seen with the gp120 loop, Tyr251 and Glu283 form a hydrogen bond and a salt bridge with maraviroc, respectively. Trp86 and Tyr89 participate in hydrophobic contacts, with Trp86 also engaging in a π-stacking interaction with the triazole group of maraviroc. A hydrogen bond with Tyr37 further anchors the triazole, while its isopropyl substituent engages in hydrophobic interactions with Met287 and Glu283. PLIP identifies three π-stacking interactions between the phenyl group of the drug and CCR5 residues Tyr108, Phe109, and Phe112. Additionally, the phenyl group forms hydrophobic contacts with Trp248, Tyr251, and Glu283. A hydrogen bond between Thr259 and a fluorine atom, as well as two hydrophobic interactions with Ile198 and Leu255, anchor the cyclohexane ring in the binding site.

Overall, maraviroc binds only a small part of the CCR5/gp120 interface, as it does not interact with the first binding site and engages only a portion of the second. This exemplifies how protein-protein interfaces are typically larger than protein-drug interfaces, making it challenging to identify a well-defined binding site for targeting protein-protein interactions with small molecules (32968059).

### BRD/H4 biological background

The bromodomain and extra-terminal (BET) family of proteins, including BRD2, BRD3, BRD4, and BRDT, play a crucial role in recognizing acetylated lysine residues on histones, particularly histone H4. This interaction plays a key role in transcriptional regulation and is implicated in various diseases, including cancer and inflammatory disorders (24751816).

In drug discovery, BET proteins are considered promising targets for small-molecule inhibitors, which disrupt their interaction with acetylated H4 and other transcriptional regulators. BET inhibitors like apabetalone and molibresib bind to the bromodomain, preventing the bromodomain from recognizing acetylated histones, thereby altering gene expression patterns. Targeting the bromodomain-histone interface is a strategy to modulate epigenetic signaling and could lead to therapeutic applications in oncology and immune-related diseases.

### BRD/H4 interactions

Bromodomains interact with acetylated histones by recognizing acetylated lysine residues on histone tails. These bromodomains share a conserved fold consisting of a left-handed helical bundle that forms a deep, largely hydrophobic binding pocket, which specifically recognizes peptide sequences containing at least one ε-N-acetylated lysine residue (24248379).

The interaction pattern detected by PLIP between BRD2’s bromo domain BD2 and the acetylated histone H4 protein involves 10 interactions and is similar to the interaction between BRD2 BD2 and the small molecule apabetalone, which engages in 8 interactions, as both bind to the hydrophobic cavity of bromodomains (see Figure 5). Both interactions involve hydrophobic interactions with BRD2 BD2 residues Val376, Leu383, and Val435, as well as at least one hydrogen bond with Asn429. Additionally, PLIP detects a π-cation interaction between BRD2 BD2 and apabetalone at His433, whereas in the BRD2 BD2–histone H4 interaction, water bridges and hydrogen bonds are observed. Notably, residues Phe372 and Tyr428 are exclusively involved in the BRD2 BD2–histone H4 interaction, while Pro371 and Asp377 are only involved in the BRD2 BD2–apabetalone interaction.

**Figure 5:**
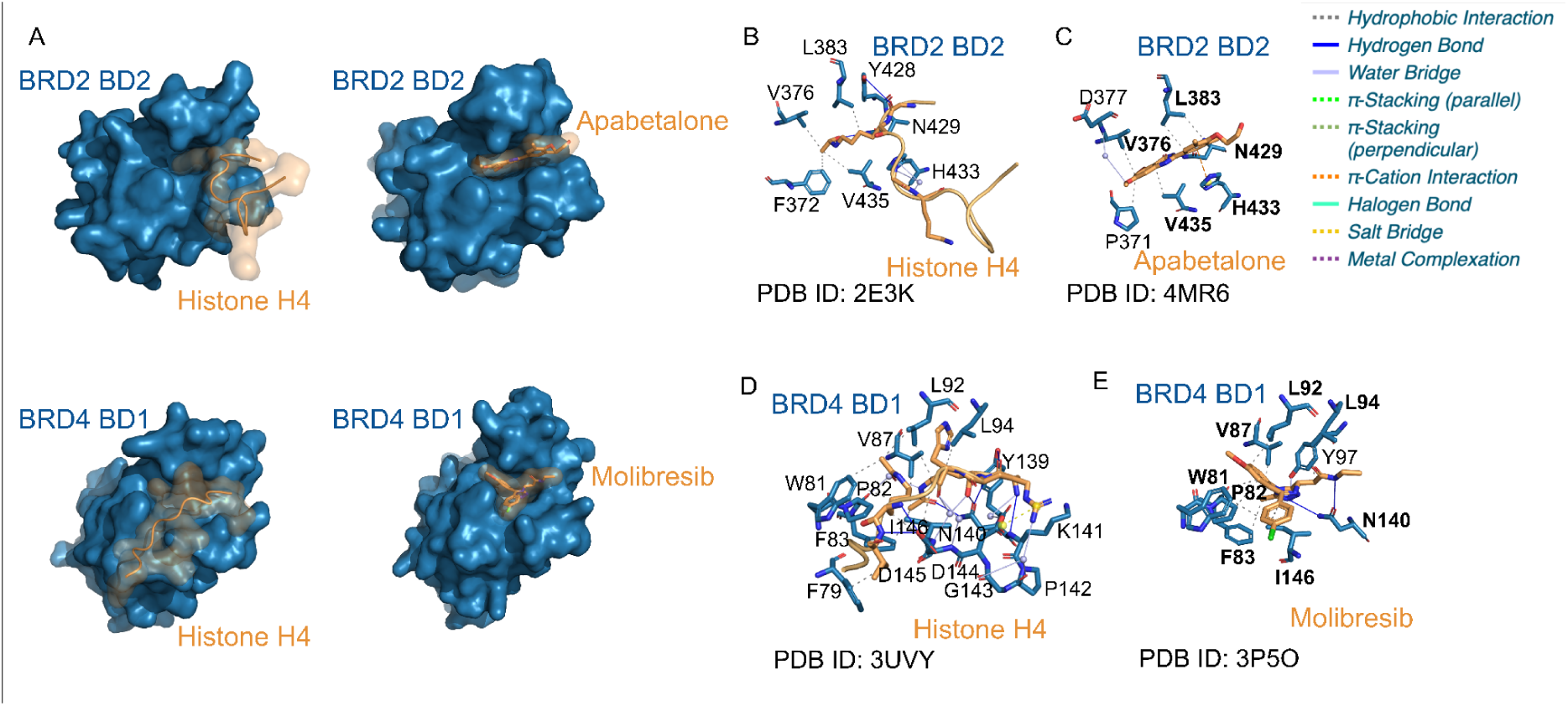
PLIP interaction pattern of the non-covalent interactions of the PPI between BRD2 BD2 and the histone H4, the PLI of BRD2 BD2 and the inhibitor apabetalone, PPI between BRD4 BD1 and histone H4 as well as the PLI between BRD4 BD1 and molibresib. **A** Surface representation of the complex structures of bromodomain (upper row BRD2 BD2, lower row BRD4 BD1, blue) with histone H4 (orange) and the inhibitors apabetalone (upper row, orange) and molibresib (lower row, orange). **B,C** non-covalent interactions detected by PLIP between BRD2 BD2 (blue) and the protein histone H4 (B, orange) and the inhibitor apabetalone (C, orange). **D,E** PLIP interaction patterns between BRD4 BD1 (blue) and the protein histone H4 (D, orange) and the inhibitor molibresib (E, orange).

Similarly, the interaction pattern between BRD4 BD1 and histone H4, which involves 28 interactions, and the interface between BRD4 BD1 and molibresib, which forms 11 interactions, show notable similarities. Both the histone and molibresib bind to the acetyl-lysine binding pocket. PLIP detects hydrophobic interactions in both complexes involving residues Trp81, Phe83, Val87, Leu92, and Leu94, as well as hydrogen bonds with Asn140. However, in contrast to the PLI, the PPI interaction between BRD4 BD1 and histone H4 involves an additional interface spanning residues Lys141 to Asp145. Specifically, PLIP detects hydrogen bonds at Lys141 and Asp145, water bridges at Pro142 and Gly143, and a salt bridge at Asp144. PLIP detects water bridges at the residues Pro82 and Ile146 for BRD4 BD1/histone and hydrophobic interactions on the same residues for BRD4 BD1/molibresib.

In both PPIs and PLIs, the interacting interfaces are relatively small and resemble a cavity. Notable similarities were observed between the respective PPIs and PLIs, particularly in terms of hydrophobic interactions and hydrogen bonding.

### Summary of examples

All of the analyzed examples show considerable overlaps in the interaction patterns between the protein-protein and protein-ligand interactions. Inhibitors are often designed to mimic the protein-protein interactions (26119925). Thus, the observed agreement in interaction patterns aligns with the design principles of these drugs. Our findings highlight that the direct displacement of the non-covalent interactions important for protein-protein binding by a small molecule is a crucial mechanism for protein interaction inhibition. Furthermore, we found a good overlap in interaction patterns among inhibitors targeting the same interface, reflecting that the development of new inhibitors is often inspired by scaffolds known to successfully inhibit the protein-protein interaction. For instance, the approved inhibitor venetoclax served as a chemical starting point for the discovery of sonrotoclax (38695063).

Commonalities in the interaction patterns of protein-protein and protein-ligand interfaces could be indicative of hot spots. In the MDM2/p53 interface, alanine scanning mutagenesis has identified the p53 residues Phe19, Leu22, and Trp23 as most important for binding (8058315). Five out of seven MDM2 residues interacting with these hot spots (Leu57, Ile61, Tyr67, Gln72, His73, Val75, Val93) are also contacted by at least one of the analyzed inhibitors (Leu57, Ile61, Tyr67, His73, Val93). The comparison of non-covalent interactions could help to identify the most important residues and interactions for binding and thereby guide drug discovery approaches.

An important consideration in the analysis of non-covalent interactions in protein complexes is the dynamics of interfaces. 3D structures are only a snapshot of the interfaces, which are subject to continuous conformational changes. In particular, the orientation of side chains is often flexible. Some interactions detected by PLIP might be lost in another conformation of the same interface while new interactions might emerge. The interaction patterns should, therefore, not be viewed as absolute.

Taken together, the comparison of PLIP profiles of protein-protein and protein-ligand interactions suggests that despite differences in size and shape of PPI and PLI, key residues and interactions are common to both. This raises the hope that it is possible in the future to automatically screen for inhibitors targeting PPI.

A premise for the success of such a screening approach is the availability of sufficient structural data. The above examples are all taken from the PDB. They are thus reliable and of high quality. But PDB is very limited in size. Therefore, the question arises whether it is possible to determine the commonalities of PPI and PLI with predicted instead of experimentally determined structures. Recent advances in structure prediction have generated tools to predict not only high-quality monomer protein structures but also PPI and PLI (34265844, 38718835, ref_protenix). However, the PPI and PLI are predicted at a lower quality than monomeric protein structures. Next, we evaluate whether such predictions are sufficient to screen for inhibitors of PPI.

### Structure prediction for PPI

The examples above used solely experimentally determined structures. How do predicted PPI and PLI compare? After the initial breakthrough in structure prediction of single proteins (34265844), AlphaFold3 (38718835) expanded capabilities by including the prediction of PPI and PLI. However, the latter is not freely available. Recently, Protenix (ref_protenix), a re-implementation of AlphaFold3, was released, which makes PLI prediction freely available. To test the quality of predicted PPI, we submitted the corresponding sequences to the AlphaFold3 and Protenix servers (see Table 4). AlphaFold3 and Protenix report the confidence in their prediction as pLDDT (Predicted Local Distance Difference Test) scores, where a score of >70% represents high and >90% very high confidence. Apart from one exception, all predictions are of very high or high confidence. Only the interaction of the receptor and HIV spike protein falls short. Since there were experimental structures for all of the examples, we could assess the actual prediction quality. The agreement of two protein structures can be assessed by the root mean square deviation (RMSD). The former assigns identical structures a value of 0. A value of less than 1 represents very good agreement. Consistent with the confidence scores, all but one PPI achieved very good RMSDs.

**Table 4:** Quality of PPI predictions. Confidence of predicted PPI (pLDDT) and root mean square deviation (RMSD) of predicted PPI against experimentally determined PPI. Confidence >70% is high, >90% very high. RMSD of <1 means structures are nearly identical. With one exception, all confidence are high or very high and RMSDs very good.

|  |  | MDM2/p53 | Bcl-2/BAX | CCR5/gp120 | XIAP/Casp9 | BRD2 BD2/H4 | BRD4 BD1/H4 |
| --- | --- | --- | --- | --- | --- | --- | --- |
| AlphaFold3 | RMSD | 0,3 | 0,7 | 21,1 | 0,7 | 0,3 | 0,4 |
|  | Confidence | 93 | 78 | 68 | 80 | 91 | 90 |
| Protenix | RMSD | 0,5 | 0,5 | 3,8 | 0,3 | 0,3 | 0,4 |
|  | Confidence | 97 | 84 | 78 | 93 | 97 | 97 |

Regarding the similarity of non-covalent interactions between experimental and predicted PPI, the results varied considerably across the five interfaces (supplementary table). Both AlphaFold and Protenix achieved excellent performance in predicting the MDM2/p53 interface, with an 82% overlap in non-covalent interactions. For the BCL-BAX interface, Protenix outperformed AlphaFold, achieving a 79% overlap compared to AlphaFold’s 69%. However, for the remaining interfaces, the agreement between predictions and reference interactions differed substantially. In the XIAP/caspase-9 complex, Protenix maintained a high overlap (79%), whereas AlphaFold’s accuracy dropped to 42%. A similar trend was observed for CCR5/gp120, where Protenix achieved 78% agreement, while AlphaFold reached only 26%. The differences were even more pronounced for the BRD-H4 interactions: in BRD2-H4, AlphaFold and Protenix showed 43% and 57% agreement, respectively, while for BRD4-H4, their overlap was considerably lower at 27% and 34%. These results indicate that while both strategies perform well for some interfaces, their predictive accuracy varies depending on the nature of the protein-protein interaction, with Protenix generally showing higher consistency across different complexes.

### Structure prediction for PLI

To compare experimental with predicted PLIs, we pursued two approaches: First, we ran structure prediction of drug and target with the Protenix server. Second, we employed docking with the state-of-the-art docking tool AutoDock using AlphaFold3 structure. Both approaches differ. The former requires only the protein sequence and a smile string of the drug. It generates a structure of their complex, i.e., the pose of the ligand, but it does not predict any affinity score for the prediction. In contrast, docking requires a structure of the protein and of the ligand, as well as a bounding box around the assumed binding site. Docking generates not only the pose of the ligand but also an affinity score. A hurdle to using docking is the definition of the bounding box. Here, we combined both approaches and derived the bounding box from the predicted PPI (see methods). This means that the docking approach is dependent on structure prediction for the placement of the bounding box.

As a ground truth and reference point for both approaches, structure prediction and docking, we used the PLIP interactions of the experimental PLI. We report the percent of molecular interactions recovered by the two approaches. We proceed in discussing the examples from best to worst.

For the MDM2 inhibitors NVP-CGM097 and SAR405838, structure prediction recovers 75 and 71% of the experimental PLIP interactions. Docking achieves 62 and 36%. For the BRD inhibitors Apabetalone and Moliresib, we obtained 71 and 89% for structure prediction and 71 and 77% for docking. In contrast, repredicting Bcl-2 inhibitor interactions resulted in very low agreement, despite the interface being predicted with high confidence in the protein-protein case. For Venetoclax, structure prediction performed worse than docking (18 vs. 36%). Similarly, for Sonrotoclax, both methods resulted in low agreement (23% each). For XIAP inhibitors, structure prediction reached 67% for both inhibitors, while docking resulted in 34% for Birinipant and 0% for Tolinapant. Finally, for the receptor CCR5, both structure prediction and docking perform not so well with 31 and 8%.

Taken together, structure prediction appears to perform better than docking. For some of the examples, there is very strong agreement with the ground-truth interactions, but for some, there is not.

### Automated prediction of novel drugs targeting novel PPIs

The performance of models like AlphaFold 3 (AF3) is often assessed using the root-mean-square deviation (RMSD) between predicted and experimentally determined Cα atoms, ideally within 2Å. While this metric provides a reliable measure of overall structural similarity, it becomes insufficient when evaluating binding interfaces. Small deviations at the atomic level can significantly affect the detection of non-covalent interactions. For experimentally resolved structures with a resolution of 2Å, it is generally assumed that interface regions in a bound state remain relatively stable and provide an accurate representation of reality. However, in the context of protein-ligand interaction prediction, it remains unclear whether this assumption holds.

Our analysis revealed a highly variable ability to repredict non-covalent interactions, depending on the protein-protein or protein-ligand system. In some cases, such as MDM2 and Bromodomain inhibitors, both structure prediction and docking achieved high agreement with experimentally determined interactions, confirming that these methods can effectively capture key interface features. However, for BCL, XIAP, and CCR5 inhibitors, the overlap was significantly lower, with some interactions missing entirely. These discrepancies highlight the challenges in accurately modeling binding interfaces, particularly when protein flexibility plays a major role.

Despite these variations, the overall interface topology was often well predicted, even when individual non-covalent interactions were not fully recovered. This suggests that while structure prediction and docking can provide valuable insights, their accuracy in modeling binding interfaces remains system-dependent. Tools like PLIP offer a crucial perspective on these predictions, helping to identify both strengths and limitations in capturing non-covalent interactions at binding sites.

## 3. Methods

UniProt accession numbers: Q00987 P04637 P10415 Q07813 P98170 P55211 Q15116 P51681 Q70145 P05107 P20701 P05362 P35222 Q92793 P25440 Q15059 O60885 P68431 P62805 P02309.

PDB ids: 4hfz 4zyf 5trf 2xa0 6o0k 8hog 1nw9 4kmp 5oqw 6met 4mbs 2e3k 4mr6 3uvy 3p5o PDB ids with chain: 4zyfA 5trfA 2xa0AC 6o0kA 8hogA 1nw9AB 4kmpA,B 5oqwB 6metBG 4mbsB 2e3kDR 4mr6A 3uvyAB 3p5oA were used.

PyMOL was used to compute buried surface area, for visualisation, and alignment using the align method.

Alphafold3. Fasta sequences from PDB files were submitted to Alphafold3 (alphafoldserver.com).

Protenix. Fasta sequences from PDB files and Smile strings from PDB (CACTVS) were submitted to Protenix (protenix-server.com).

PLIP (33950214) was used to assess molecular interactions. For Table 1, water bridge distances from the two interaction partners to water were summed up.

PLIP statistics for PPI were computed for PDB ids in SCOP2 (31724711), which have exactly two chains A and B (file scop-cla-latest.txt).

PLI statistics for PLI were computed for all PDB ids, which have a bound ligand. Ions are excluded from the analysis.

AutoDock. GNINA (github.com/gnina/gnina), a fork of AutoDock Vina, was used for docking. Ligands downloaded from the PDB were parsed as .sdf files. The bounding box required for AutoDock was derived from the PPI interfaces of predicted structures (AlphaFold3) as detected by PLIP. For the atoms in the PLIP interactions, the minimal and maximal *x, y, z* values were computed. Using an additional margin of 8Å, the bounding box is defined by *(x_min_,y_min_,z_min_)-*(*8,8,8*) to *(x_max_,y_max_,z_max_)+*(*8,8,8*). The ligand is placed at the center of mass of the atoms of the PLIP interactions.

## 4. Discussion and conclusion

Although PPIs are challenging drug targets, they hold great promise in drug discovery because they are at the heart of many biological processes and, thus, disease mechanisms. Targeting an interface that is already involved in a PPI by a drug represents a special case of promiscuity. Commonalities between PPI and PLI could serve as a strong basis for drug screenings of PPI modulators and drug repositioning. To evaluate the similarity of protein-protein and protein-ligand interfaces, we have analyzed the non-covalent interaction patterns of 9 PPI inhibitors targeting 5 different protein-protein interfaces using PLIP.

We found that PPIs have, in general, more non-covalent interactions than PLIs, which is consistent with previous observations that protein-protein interfaces are typically larger than protein-ligand interfaces (32968059). In particular, the CCR5/gp120 and XIAP/caspase-9 interfaces are significantly bigger than the corresponding PLIs. However, in the selected examples of PPI drugs in clinical trials, all protein-protein interfaces exhibit fewer interactions than the average PPI in the PDB. The crystallization of globular domains is easier than that of peptide segments, which introduces an inherent bias towards globular proteins in the PDB. This might lead to an overrepresentation of large protein-protein interfaces between globular domains in the data. On the other hand, our findings could be indicative of small PPIs being more druggable than large ones. Hence, successful PPI inhibition tends to involve relatively small interfaces. Notably, the bromodomain/histone interface forms a deep cavity, which resembles classic ligand binding sites. Interestingly, the number of interactions formed by SAR405838, venetoclax, and sonrotoclax is above the average PLI. This suggests that drugs might require the ability to form a higher number of interactions to mimic PPIs.

Our PLIP analysis shows considerable overlaps in interfaces and non-covalent interactions between PPI and PLI. It demonstrates how PLIs bind protein-protein interfaces in a promiscuous manner by mimicking the non-covalent interaction patterns of the PPIs. This makes drug screenings for PPI modulation based on binding site characteristics and interaction patterns a promising approach for drug discovery. Furthermore, the comparison of PPIs and PLIs could be a strong foundation for drug repositioning. Nevertheless, certain considerations have to be taken into account. First, the size difference of PPIs and PLIs requires a careful evaluation of the interfaces and identification of the most important regions for binding. The concept of hot spots could be helpful in this context. Second, PLIs can use a larger set of interaction types than PPIs because the diversity of chemical moieties is higher than that of native amino acids. For instance, we found halogen bonds in the interaction patterns of SAR405838 and maraviroc. Screening approaches have to capture this difference in chemical diversity.

The limited availability of experimentally determined protein structures restricts a systematic screening of PPI inhibitors. Therefore, we wished to examine the quality of protein structure predictions for the characterization of interfaces. Re-predictions of the example complex structures demonstrated an overall good structural agreement. Yet, how well the PLIP interactions could be recovered greatly differs between the examples. Moreover, performance was dependent on the interaction prediction method. While structure prediction methods have a huge potential to expand the amount of data for drug screenings, they have to be used with care and currently still require individual evaluation. PLIP interaction patterns could facilitate the assessment of protein structure predictions for interface characterizations. Taken together, our analysis shows that PPI and PLI have sufficient commonalities to merit future work into computational screening for drugs targeting PPI. It will be key to further improve structure prediction, specifically for binding sites, and to deal with flexibility.

## Acknowledgement

We would like to thank Alex Meschiavelli for technical support and BMBF scads.ai for funding. Thanks to Sebastian Salentin for initiating and implementing PLIP.

## Contributions

SB, PS, MS conceived the study, SB, PS, CS, MS analysed data and wrote the manuscript.

---

The following PMIDs are cited in the article (in order of appearance):

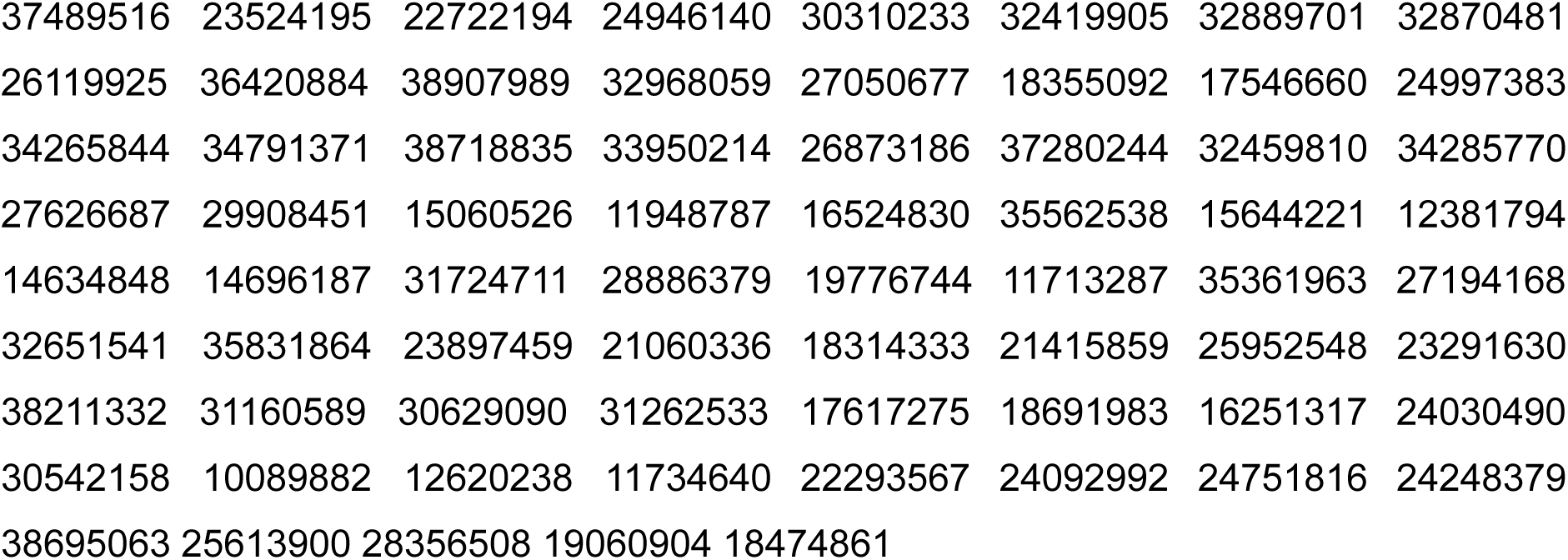

## References

1. Farzan M,…, Choe H. Tyrosine sulfation of the amino terminus of CCR5 facilitates HIV-1 entry. Cell. 1999 https://pubmed.ncbi.nlm.nih.gov/10089882

2. Lohrum MA,…, Vousden KH. C-terminal ubiquitination of p53 contributes to nuclear export. Mol Cell Biol. 2001 https://pubmed.ncbi.nlm.nih.gov/11713287

3. Renatus M,…, Salvesen GS. Dimer formation drives the activation of the cell death protease caspase 9. Proc Natl Acad Sci U S A. 2001 https://pubmed.ncbi.nlm.nih.gov/11734640

4. Chakrabarti P and Janin J. Dissecting protein-protein recognition sites. Proteins. 2002 https://pubmed.ncbi.nlm.nih.gov/11948787

5. Kortemme T and Baker D. A simple physical model for binding energy hot spots in protein-protein complexes. Proc Natl Acad Sci U S A. 2002 https://pubmed.ncbi.nlm.nih.gov/12381794

6. Shiozaki EN,…, Shi Y. Mechanism of XIAP-mediated inhibition of caspase-9. Mol Cell. 2003 https://pubmed.ncbi.nlm.nih.gov/12620238

7. Gao Y,…, Lai L. Structure-based method for analyzing protein-protein interfaces. J Mol Model. 2004 https://pubmed.ncbi.nlm.nih.gov/14634848

8. Sarkhel S and Desiraju GR. N-H…O, O-H…O, and C-H…O hydrogen bonds in protein-ligand complexes: strong and weak interactions in molecular recognition. Proteins. 2004 https://pubmed.ncbi.nlm.nih.gov/14696187

9. Keskin O,…, Nussinov R. Hot regions in protein--protein interactions: the organization and contribution of structurally conserved hot spot residues. J Mol Biol. 2005 https://pubmed.ncbi.nlm.nih.gov/15644221

10. Dorr P,…, Perros M. Maraviroc (UK-427,857), a potent, orally bioavailable, and selective small-molecule inhibitor of chemokine receptor CCR5 with broad-spectrum anti-human immunodeficiency virus type 1 activity. Antimicrob Agents Chemother. 2005 https://pubmed.ncbi.nlm.nih.gov/16251317

11. Blundell TL,…, Burke D. Structural biology and bioinformatics in drug design: opportunities and challenges for target identification and lead discovery. Philos Trans R Soc Lond B Biol Sci. 2006 https://pubmed.ncbi.nlm.nih.gov/16524830

12. Moreira IS,…, Ramos MJ. Hot spots--a review of the protein-protein interface determinant amino-acid residues. Proteins. 2007 https://pubmed.ncbi.nlm.nih.gov/17546660

13. Este JA and Telenti A. HIV entry inhibitors. Lancet. 2007 https://pubmed.ncbi.nlm.nih.gov/17617275

14. Chipuk JE and Green DR. How do BCL-2 proteins induce mitochondrial outer membrane permeabilization. Trends Cell Biol. 2008 https://pubmed.ncbi.nlm.nih.gov/18314333

15. Stumpf MP,…, Wiuf C. Estimating the size of the human interactome. Proc Natl Acad Sci U S A. 2008 https://pubmed.ncbi.nlm.nih.gov/18474861

16. Lieberman-Blum SS,…, Bandres JC. Maraviroc: a CCR5-receptor antagonist for the treatment of HIV-1 infection. Clin Ther. 2008 https://pubmed.ncbi.nlm.nih.gov/18691983

17. Venkatesan K,…, Vidal M. An empirical framework for binary interactome mapping. Nat Methods. 2009 https://pubmed.ncbi.nlm.nih.gov/19060904

18. Levine AJ and Oren M. The first 30 years of p53: growing ever more complex. Nat Rev Cancer. 2009 https://pubmed.ncbi.nlm.nih.gov/19776744

19. Ku B,…, Oh BH. Evidence that inhibition of BAX activation by BCL-2 involves its tight and preferential interaction with the BH3 domain of BAX. Cell Res. 2011 https://pubmed.ncbi.nlm.nih.gov/21060336

20. Kelly PN and Strasser A. The role of Bcl-2 and its pro-survival relatives in tumourigenesis and cancer therapy. Cell Death Differ. 2011 https://pubmed.ncbi.nlm.nih.gov/21415859

21. Fulda S and Vucic D. Targeting IAP proteins for therapeutic intervention in cancer. Nat Rev Drug Discov. 2012 https://pubmed.ncbi.nlm.nih.gov/22293567

22. Lounkine E,…, Urban L. Large-scale prediction and testing of drug activity on side-effect targets. Nature. 2012 https://pubmed.ncbi.nlm.nih.gov/22722194

23. Souers AJ,…, Elmore SW. ABT-199, a potent and selective BCL-2 inhibitor, achieves antitumor activity while sparing platelets. Nat Med. 2013 https://pubmed.ncbi.nlm.nih.gov/23291630

24. Hu Y and Bajorath J. Compound promiscuity: what can we learn from current data. Drug Discov Today. 2013 https://pubmed.ncbi.nlm.nih.gov/23524195

25. Anil B,…, Noble ME. The structure of an MDM2-Nutlin-3a complex solved by the use of a validated MDM2 surface-entropy reduction mutant. Acta Crystallogr D Biol Crystallogr. 2013 https://pubmed.ncbi.nlm.nih.gov/23897459

26. Tan Q,…, Wu B. Structure of the CCR5 chemokine receptor-HIV entry inhibitor maraviroc complex. Science. 2013 https://pubmed.ncbi.nlm.nih.gov/24030490

27. Dubrez L,…, Glorian V. IAP proteins as targets for drug development in oncology. Onco Targets Ther. 2013 https://pubmed.ncbi.nlm.nih.gov/24092992

28. Picaud S,…, Filippakopoulos P. RVX-208, an inhibitor of BET transcriptional regulators with selectivity for the second bromodomain. Proc Natl Acad Sci U S A. 2013 https://pubmed.ncbi.nlm.nih.gov/24248379

29. Filippakopoulos P and Knapp S. Targeting bromodomains: epigenetic readers of lysine acetylation. Nat Rev Drug Discov. 2014 https://pubmed.ncbi.nlm.nih.gov/24751816

30. Anighoro A,…, Rastelli G. Polypharmacology: challenges and opportunities in drug discovery. J Med Chem. 2014 https://pubmed.ncbi.nlm.nih.gov/24946140

31. Cukuroglu E,…, Keskin O. Hot spots in protein-protein interfaces: towards drug discovery. Prog Biophys Mol Biol. 2014 https://pubmed.ncbi.nlm.nih.gov/24997383

32. Uhlen M,…, Ponten F. Proteomics. Tissue-based map of the human proteome. Science. 2015 https://pubmed.ncbi.nlm.nih.gov/25613900

33. Delbridge AR and Strasser A. The BCL-2 protein family, BH3-mimetics and cancer therapy. Cell Death Differ. 2015 https://pubmed.ncbi.nlm.nih.gov/25952548

34. Pelay-Gimeno M,…, Grossmann TN. Structure-Based Design of Inhibitors of Protein-Protein Interactions: Mimicking Peptide Binding Epitopes. Angew Chem Int Ed Engl. 2015 https://pubmed.ncbi.nlm.nih.gov/26119925

35. Haupt VJ,…, Schroeder M. Computational Drug Repositioning by Target Hopping: A Use Case in Chagas Disease. Curr Pharm Des. 2016 https://pubmed.ncbi.nlm.nih.gov/26873186

36. Scott DE,…, Skidmore J. Small molecules, big targets: drug discovery faces the protein-protein interaction challenge. Nat Rev Drug Discov. 2016 https://pubmed.ncbi.nlm.nih.gov/27050677

37. Oliner JD,…, Caenepeel S. The Role of MDM2 Amplification and Overexpression in Tumorigenesis. Cold Spring Harb Perspect Med. 2016 https://pubmed.ncbi.nlm.nih.gov/27194168

38. Heinrich JC,…, Schroeder M. New HSP27 inhibitors efficiently suppress drug resistance development in cancer cells. Oncotarget. 2016 https://pubmed.ncbi.nlm.nih.gov/27626687

39. Finan C,…, Casas JP. The druggable genome and support for target identification and validation in drug development. Sci Transl Med. 2017 https://pubmed.ncbi.nlm.nih.gov/28356508

40. Kastenhuber ER and Lowe SW. Putting p53 in Context. Cell. 2017 https://pubmed.ncbi.nlm.nih.gov/28886379

41. Ran X and Gestwicki JE. Inhibitors of protein-protein interactions (PPIs): an analysis of scaffold choices and buried surface area. Curr Opin Chem Biol. 2018 https://pubmed.ncbi.nlm.nih.gov/29908451

42. Pushpakom S,…, Pirmohamed M. Drug repurposing: progress, challenges and recommendations. Nat Rev Drug Discov. 2019 https://pubmed.ncbi.nlm.nih.gov/30310233

43. Shaik MM,…, Chen B. Structural basis of coreceptor recognition by HIV-1 envelope spike. Nature. 2019 https://pubmed.ncbi.nlm.nih.gov/30542158

44. Birkinshaw RW,…, Czabotar PE. Structures of BCL-2 in complex with venetoclax reveal the molecular basis of resistance mutations. Nat Commun. 2019 https://pubmed.ncbi.nlm.nih.gov/31160589

45. . Chen B. Molecular Mechanism of HIV-1 Entry. Trends Microbiol. 2019 https://pubmed.ncbi.nlm.nih.gov/31262533

46. Andreeva A,…, Murzin AG. The SCOP database in 2020: expanded classification of representative family and superfamily domains of known protein structures. Nucleic Acids Res. 2020 https://pubmed.ncbi.nlm.nih.gov/31724711

47. Parisi D,…, Schroeder M. Drug repositioning or target repositioning: A structural perspective of drug-target-indication relationship for available repurposed drugs. Comput Struct Biotechnol J. 2020 https://pubmed.ncbi.nlm.nih.gov/32419905

48. Adasme MF,…, Schroeder M. Structure-based drug repositioning explains ibrutinib as VEGFR2 inhibitor. PLoS One. 2020 https://pubmed.ncbi.nlm.nih.gov/32459810

49. Konopleva M,…, Andreeff M. MDM2 inhibition: an important step forward in cancer therapy. Leukemia. 2020 https://pubmed.ncbi.nlm.nih.gov/32651541

50. Lamb YN. Remdesivir: First Approval. Drugs. 2020 https://pubmed.ncbi.nlm.nih.gov/32870481

51. Singh TU,…, Singh RK. Drug repurposing approach to fight COVID-19. Pharmacol Rep. 2020 https://pubmed.ncbi.nlm.nih.gov/32889701

52. Lu H,…, Shi J. Recent advances in the development of protein-protein interactions modulators: mechanisms and clinical trials. Signal Transduct Target Ther. 2020 https://pubmed.ncbi.nlm.nih.gov/32968059

53. Adasme MF,…, Schroeder M. PLIP 2021: expanding the scope of the protein-ligand interaction profiler to DNA and RNA. Nucleic Acids Res. 2021 https://pubmed.ncbi.nlm.nih.gov/33950214

54. Jumper J,…, Hassabis D. Highly accurate protein structure prediction with AlphaFold. Nature. 2021 https://pubmed.ncbi.nlm.nih.gov/34265844

55. Bolz SN,…, Schroeder M. Structural binding site comparisons reveal Crizotinib as a novel LRRK2 inhibitor. Comput Struct Biotechnol J. 2021 https://pubmed.ncbi.nlm.nih.gov/34285770

56 Varadi M,…, Velankar S. AlphaFold Protein Structure Database: massively expanding the structural coverage of protein-sequence space with high-accuracy models. Nucleic Acids Res. 2022 https://pubmed.ncbi.nlm.nih.gov/34791371

57. Kennedy MC and Lowe SW. Mutant p53: it’s not all one and the same. Cell Death Differ. 2022 https://pubmed.ncbi.nlm.nih.gov/35361963

58. Alzyoud L,…, Ghattas MA. Structure-based assessment and druggability classification of protein-protein interaction sites. Sci Rep. 2022 https://pubmed.ncbi.nlm.nih.gov/35562538

59. Zhu H,…, Zhou Z. Targeting p53-MDM2 interaction by small-molecule inhibitors: learning from MDM2 inhibitors in clinical trials. J Hematol Oncol. 2022 https://pubmed.ncbi.nlm.nih.gov/35831864

60. Burley SK,…, Zardecki C. RCSB Protein Data Bank (RCSB.org): delivery of experimentally-determined PDB structures alongside one million computed structure models of proteins from artificial intelligence/machine learning. Nucleic Acids Res. 2023 https://pubmed.ncbi.nlm.nih.gov/36420884

61. Schake P,…, Schroeder M. An interaction-based drug discovery screen explains known SARS-CoV-2 inhibitors and predicts new compound scaffolds. Sci Rep. 2023 https://pubmed.ncbi.nlm.nih.gov/37280244

62. Bolz SN and Schroeder M. Promiscuity in drug discovery on the verge of the structural revolution: recent advances and future chances. Expert Opin Drug Discov. 2023 https://pubmed.ncbi.nlm.nih.gov/37489516

63. Liu J,…, Liu Y. Sonrotoclax overcomes BCL2 G101V mutation-induced venetoclax resistance in preclinical models of hematologic malignancy. Blood. 2024 https://pubmed.ncbi.nlm.nih.gov/38211332

64. Guo Y,…, Wang Z. Discovery of the Clinical Candidate Sonrotoclax (BGB-11417), a Highly Potent and Selective Inhibitor for Both WT and G101V Mutant Bcl-2. J Med Chem. 2024 https://pubmed.ncbi.nlm.nih.gov/38695063

65. Abramson J,…, Jumper JM. Accurate structure prediction of biomolecular interactions with AlphaFold 3. Nature. 2024 https://pubmed.ncbi.nlm.nih.gov/38718835

66. Cankara F,…, Keskin O. DiPPI: A Curated Data Set for Drug-like Molecules in Protein-Protein Interfaces. J Chem Inf Model. 2024 https://pubmed.ncbi.nlm.nih.gov/38907989

67. REF_protenix. Chen X,…, Xiao W. Protenix - Advancing Structure Prediction Through a Comprehensive AlphaFold3 Reproduction. bioRxiv. 2025 https://www.biorxiv.org/content/10.1101/2025.01.08.631967v1

