## Supplementary figures and images for "The structural basis of drugs targeting protein-protein interactions uncovered with the protein-ligand interaction profiler PLIP"

### supplementary table

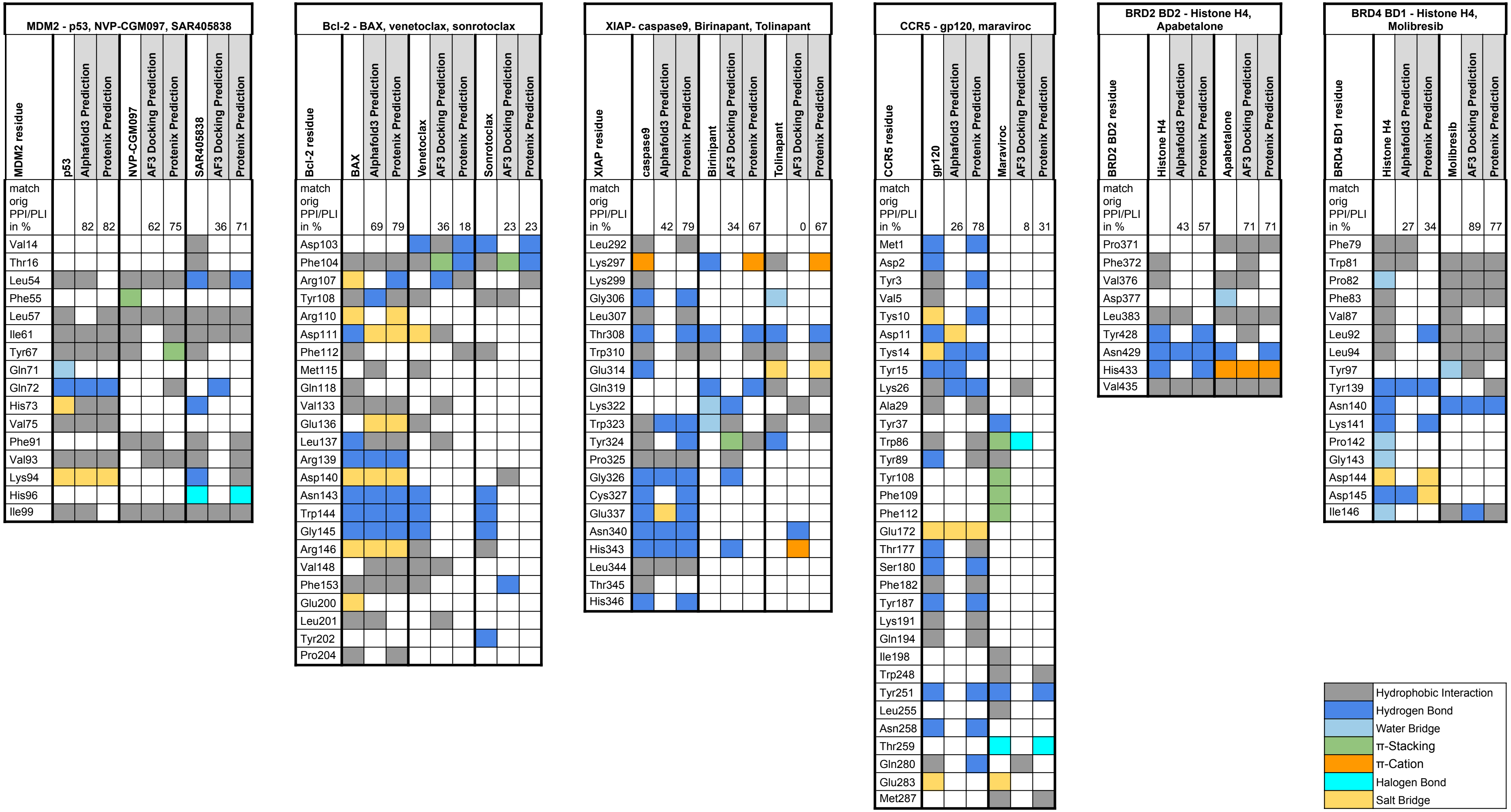
